# Reflux of Endoplasmic Reticulum proteins to the cytosol yields inactivation of tumor suppressors

**DOI:** 10.1101/2020.04.13.038935

**Authors:** Daria Sicari, Raphael Pineau, Pierre-Jean Le Reste, Luc Negroni, Sophie Chat, Aiman Mohtar, Daniel Thomas, Reynald Gillet, M. Ted Hupp, Eric Chevet, Aeid Igbaria

**Affiliations:** Inserm U1242, University of Rennes, Rennes, France; Centre de lutte contre le cancer Eugène Marquis, Rennes, France; Neurosurgery Dept, University Hospital of Rennes, 35000 Rennes, France; Institut de Génétique et de Biologie Moléculaire et Cellulaire, 67404 Illkirch, France; Centre National de la Recherche Scientifique, UMR7104, 67404 Illkirch, France; Institut National de la Santé et de la Recherche Médicale, U1258, 67404 Illkirch, France; Université de Strasbourg, 67404 Illkirch, France; Univ. Rennes, CNRS, Institut de Génétique et Développement de Rennes (IGDR) UMR6290, 35000 Rennes, France; Edinburgh Cancer Research Centre at the Institute of Genetics and Molecular Medicine, Edinburgh University, Edinburgh, UK; International Centre for Cancer Vaccine Science, Gdansk, Poland; Department of Life Sciences, Ben-Gurion University of the Negev, Beer Sheva 8410501, Israel

## Abstract

In the past decades many studies reported Endoplasmic Reticulum (ER) resident proteins to localize to the cytosol but the mechanisms by which this occurs and whether these proteins exert cytosolic functions remain unknown. We found that select ER luminal proteins accumulate in the cytosol of glioblastoma cells isolated from mouse and human tumors. In cultured cells ER protein reflux to the cytosol occurs upon proteostasis perturbation. As such we investigated whether refluxed proteins gain new functions in the cytosol thus providing advantage to tumor cells. Using the ER luminal protein AGR2 as a model, we showed that it is refluxed to the cytosol where it binds and inhibits the tumor suppressor p53. We named this phenomenon ER to Cytosol Signaling (ERCYS) as an ER surveillance mechanism conserved in Eukaryotes to relieve the ER from its contents upon stress and to provide selective advantage to tumor cells through gain-of-cytosolic functions.

## INTRODUCTION

The Endoplasmic Reticulum (ER) is the gateway to the secretory pathway thus maintaining the communication between the cell’s intracellular and extracellular environment. In addition, the ER is a sensing organelle that coordinates many stress signaling pathways (Higa and Chevet 2012; Hetz, Chevet, and Oakes 2015; Alexia et al. 2013). Secretory and transmembrane proteins translocate into the ER through the Sec61 translocon. The ER is crowded with molecular chaperones and foldases that ensure these proteins’ productive folding followed by their export en-route to their final destination (Rapoport 2007). Diverse perturbations compromise the folding and maturation of secretory proteins in the ER thereby causing ER stress. To ensure productive folding, cells have also evolved various ER quality control (ERQC) systems allowing for further folding rounds (Adams, Oster, and Hebert 2019) and their degradation in the cytosol by a process termed ER-associated degradation (ERAD) (Rutkowski et al. 2006; Travers et al. 2000; Vembar and Brodsky 2008). In addition to ERQC and ERAD, a pre-emptive quality control (pre-QC) mechanism was also described that averts protein entry into the secretory pathway under protein-folding stress resulting in their proteasomal degradation in the cytosol (Kang et al. 2006). If these quality control systems are overwhelmed, ER stress activates a signaling pathway called the Unfolded Protein Response (UPR) that augments the ER protein folding capacity through transcriptional upregulation of genes encoding ER chaperones, oxidoreductases, lipid biosynthetic enzymes and ERAD components. The UPR is transduced by three transmembrane sensor proteins (PERK, ATF6α and IRE1α) that sense and monitor the protein folding status of the ER through their luminal domains and transmit signals to the rest of the cell through their cytosolic domain (Almanza et al. 2019).

In the past three decades a subset of ER resident proteins were reported to accumulate in the cytosol of several human diseases including cancer and neurodegenerative diseases (Afshar, Black, and Paschal 2005; Galligan and Petersen 2012; Shim et al. 2018; Tarr et al. 2010; Turano et al. 2002; Wiersma et al. 2015). For instance, the localization of ER-resident proteins to different cellular compartments has been extensively reported including members of the Protein Disulfide Isomerase (PDI) family, GRP78/BiP, calreticulin and others (Afshar, Black, and Paschal 2005; Galligan and Petersen 2012; Shim et al. 2018; Tarr et al. 2010; Turano et al. 2002; Wiersma et al. 2015). Despite this recurring observation, the potential functions of those proteins in the cytosol remain unclear.

Recently, we showed that protein folding stress causes ER resident proteins to be refluxed to the cytosol in the yeast *Saccharomyces Cerevisiae* (Igbaria et al. 2019). This mechanism requires ER and cytosolic chaperones and co-chaperones but is independent of the ERAD machinery and protein degradation (Igbaria et al. 2019). Here we found that ER stress mediated protein reflux is conserved in mammalian cells and in isolated cancer cells from human and murine tumors and aims at debulking the ER upon stress. Moreover, we found that this process may be constitutively active in tumor cells and lead to cytosolic gain-of-functions of the refluxed protein as inhibitor of tumor suppressors, thereby exhibiting pro-oncogenic features.

## RESULTS

### ER resident proteins are refluxed from the ER lumen to the cytosol in cancer cells isolated from human and murine GBM tumors

To study the role of ER protein reflux in tumors, we focused on Glioblastoma multiforme (GBM) in which the Unfolded Protein Response (UPR) sustains tumor aggressiveness (Obacz et al. 2017). Mouse GBM cells (GL261) were grafted orthotopically in the brain of C57BL/6 mice and 30 days post-injection, tumors were isolated, dissociated and isolated tumor cells subjected to subcellular fractionation using previously validated fractionation methods (Holden and Horton 2009). Immunoblot analysis of the digitonin fraction (enriched in cytosolic proteins) from freshly isolated tumor cells was compared to that of dissociated control tissue from the opposite hemisphere of the brain (non-tumor). It revealed higher levels of ER-resident proteins in tumor cells’ cytosol than in the non-tumor counterpart (Figure 1A–1F and Figure S1A-B). Some ER-resident proteins were enriched up to ~70% in the cytosolic fraction compared to only ~10% enrichment in non-tumor controls. These results indicate that tumor cells are more prone to exhibit reflux of ER proteins to the cytosol than non-tumor cells. We confirmed these findings in another tumor model. Namely, U87 cells orthotopically implanted in NSG mouse brains (Figure S1C). In both GBM models (GL261 and U87), N-linked-glycoproteins (such as ERDJ3/DNAJB11) were found in the digitonin fraction thus indicating that the refluxed proteins had been translocated into the ER and modified by N-Linked glycosylation, before being refluxed to the cytosol.

**Figure 1:**
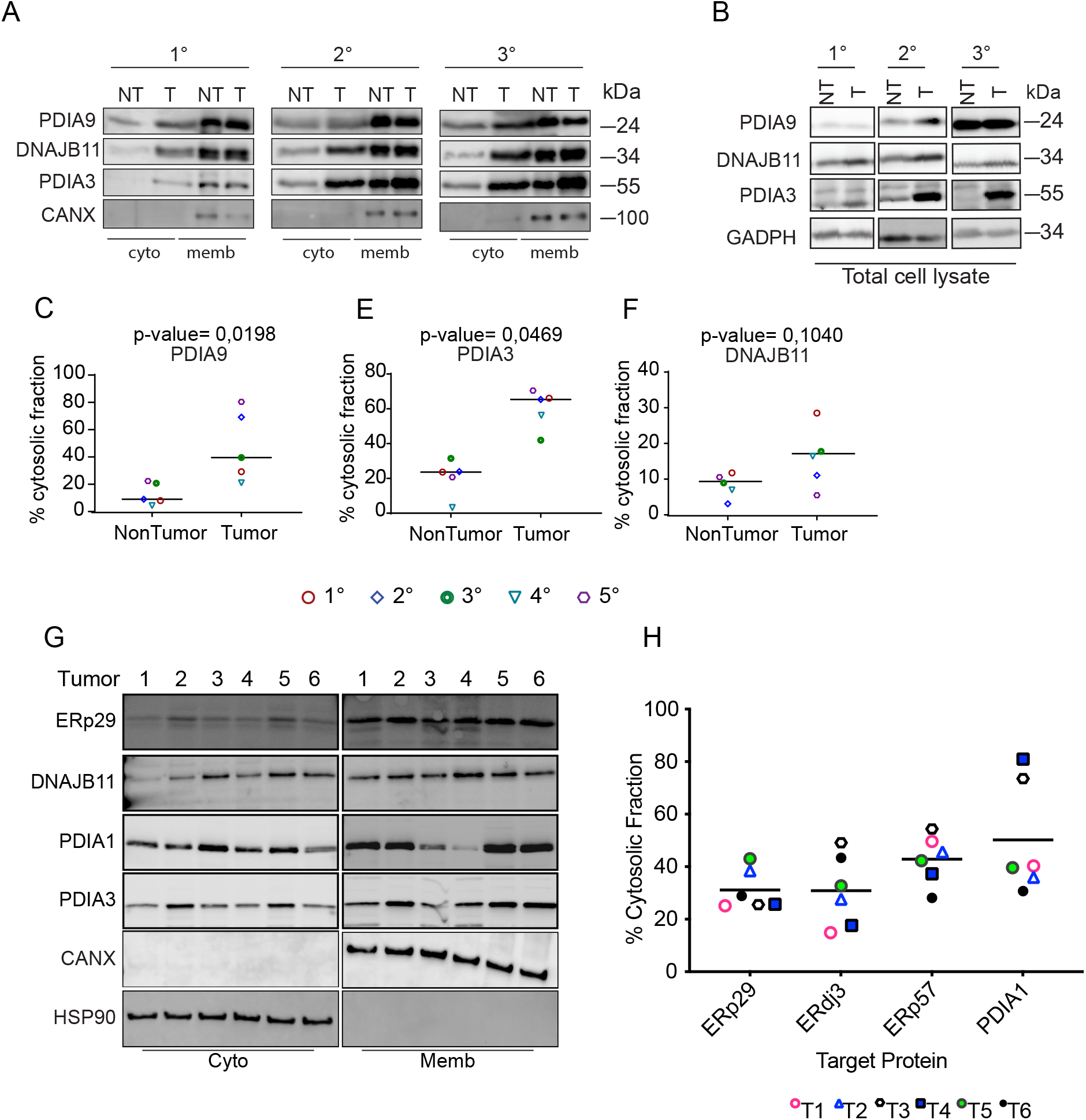
ER proteins are rerouted to the cytosol in human and mouse Glioblastoma (GBM) tumors. (A) Representative western blots after subcellular protein fractionation experiment in isolated murine-derived non-tumor (NT) and GBM tumor (T) tissues. (B) Total levels of the ER-lumenal proteins tested from the cell lysate derived from tissues used in (A). (C-F) Quantification of the protein levels of ER lumenal proteins in the cytosolic fraction as shown in (A). (G-H) Human derived GBM tumors were processed as in A. Representative western blot were performed (G) and the percentage of ER-lumenal protein cytosolic localization (H) were quantified. Data are the average from six different tumors.

We next asked whether this phenomenon is also observed in human GBM samples freshly isolated from patients at surgery (Table S1). We tested the subcellular protein localization of ER resident proteins in freshly isolated human tumor cells. Tumor tissues were dissociated, and digitonin fractions tested for the presence of ER luminal proteins using immunoblotting. In the majority of tumors (80% of the tested tumors) ~50% of the ER proteins evaluated were detected in the digitonin fraction including ERDJ3/DNAJB11 in its N-glycosylated state (Figure 1G-H and Figure S1D). Moreover, individual tumors exhibited heterogenous refluxed protein patterns, which might reflect inter-tumor heterogeneity (Figure 1G and 1H). These data indicate that in these tumors, ER protein reflux might be selectively regulated by different factors such as tumor heterogeneity, genetic background or activation status of UPR. These findings will stimulate others to replicate and extend these data in other tumor models.

### ER stress mediates ER-resident proteins reflux from the ER to the cytosol

We next sought to identify factors regulating ER-to-cytosol protein reflux. Recently, we reported that ER luminal proteins were refluxed to the cytosol upon ER stress in the yeast *S. cerevisiae* (Igbaria et al. 2019; Lajoie and Snapp 2020) in a chaperone-mediated process (Igbaria et al. 2019). We tested whether ER stress/UPR activation – two factors that showed a correlation with protein reflux in *S. cerevisiae* – would also cause ER protein reflux in mammalian cells. As such we monitored the localization of an engineered ER targeted super-folder GFP (ER-sfGFP) in HEK293T cells using confocal microscopy, to follow the fate of ER-sfGFP in living cells. Notably, HEK293T cells treated with Tunicamycin (Tm), which perturbs protein folding by inhibiting N-linked glycosylation, showed enhanced cytosolic localization of ER-sfGFP, this localization reached a maximum after 24hrs of treatment (Figure 2A). To confirm that ER-resident proteins found in the cytosol after ER stress originated from the ER lumen, we engineered an ER targeted photoactivatable fluorescent protein (FP) called mEOS3.2 by adding ER signal peptide and a KDEL retention sequence, that discriminates newly synthesized proteins from the pre-existing pool. Indeed, UV exposure shifts the excitation maxima of the mEOS3.2 from 488nm to 573nm, allowing detection of proteins synthesized before a UV pulse exposure (Figure S2A). Notably, after a UV pulse and Tm or DMSO treatments for 24 hours, the mEOS3.2^573^ pool was mainly localized in the ER of the DMSO treated cells, but cells treated with Tm or the ER-Golgi transport inhibitor Brefeldin A (BFA) showed a significant fraction of mEOS3.2^573^ localized in the cytosol (Figure 2B). These results indicated that during stress pre-existing and ER localized proteins are refluxed to the cytosol where they might exist in a folded, functional state.

**Figure 2:**
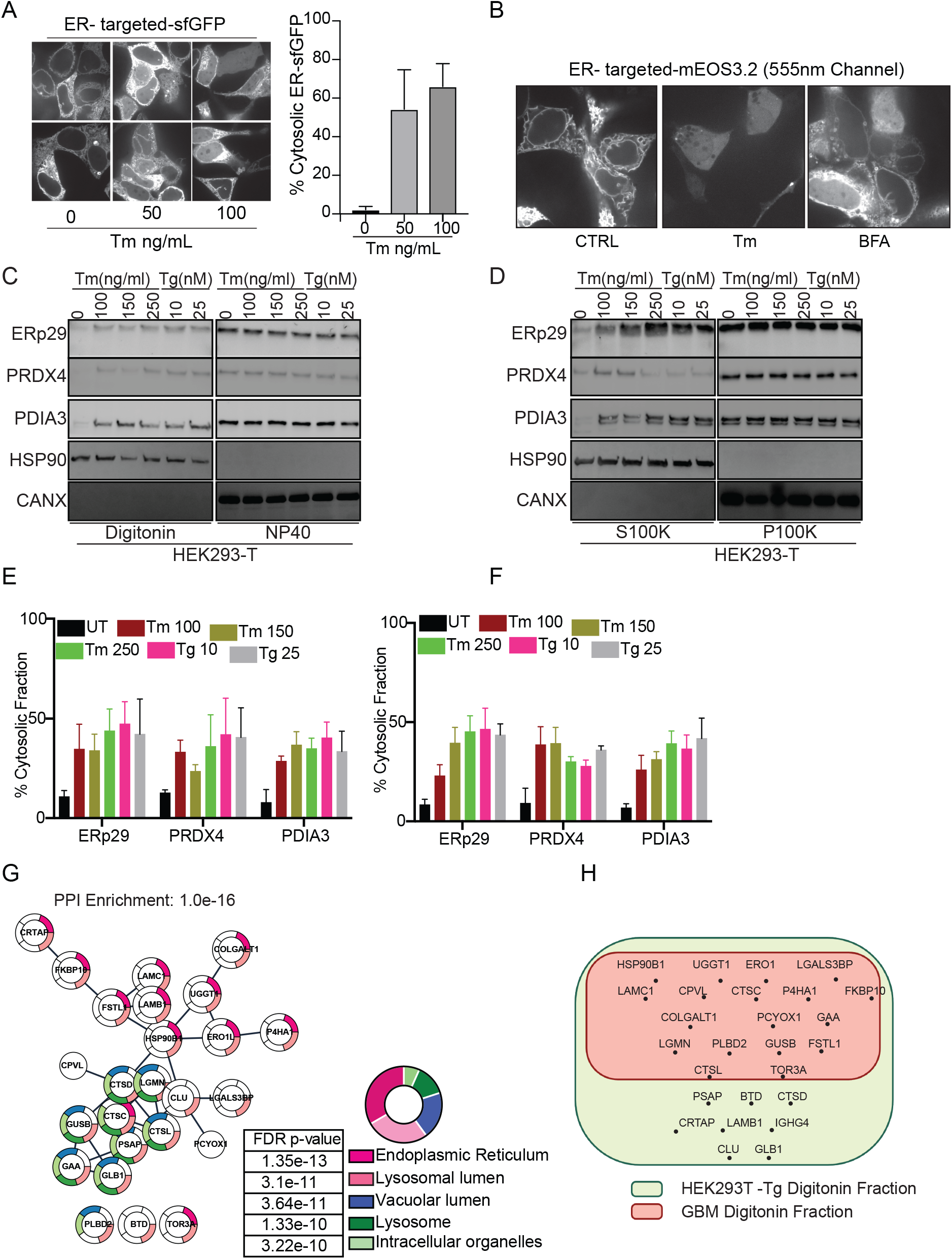
pre-existing, ER targeted mEOS3.2 and ER endogenous proteins reflux to the cytosol during ER stress. (A) HEK293T were transfected with super-folder GFP and treated with Tm at the indicated concentrations for 16 hours. Graph shows the quantification of three independent experiments (B) Microscopy images of HEK293T cells expressing ER-targeted mEos3.2, UV light was used to first convert the mEos3.2 then images were taken at 550 nm in the absence or presence of Tm and BFA. (C,D) Subcellular protein fractionation of several ER resident proteins in HEK293T cells treated with the indicated concentrations of Tm or Tg for 16 hours using Digitonin (C) or differential centrifugation (D), representative western blots are showed. (E,F) Quantification of the subcellular protein fractionation of several ER endogenous proteins in HEK293T cells treated with different concentrations of Tm and Tg for 16 hrs from panels C and D respectively. (G) Mass spec analysis of soluble ER targeted glycoproteins in HEK293T cells treated with Tg. (H) Mass spectrometry analysis of cytosolically located soluble ER glycoproteins in HEK293T cells treated with Tg compared to that found in human GBM tumor cells-derived cytosols.

We next investigated whether endogenous ER-resident proteins were also refluxed to the cytosol in cultured cells that were exposed to various ER stress inducers. Subcellular protein fractionation using minimal concentration of digitonin that results in proper isolation of the different subcellular fractions was carried out in cells subjected to ER stress induced by Tm or Thapsigargin (Tg) which inhibits the sarco-endoplasmic reticulum Ca^2+^ ATPase. This was followed by an analysis of the localization of different endogenous ER resident proteins including the soluble ERp29/PDIA9, PRDX4, ERp57/PDIA3, and the integral protein calnexin (CANX). We found that soluble ER luminal proteins were enriched in the digitonin fraction up to 50-55% (Figure 2C and 2E) but not calnexin, thus indicating that ER reflux could be exclusive for soluble proteins. To rule out the possibility that ER stress may change the composition of the ER membrane thereby sensitizing it to digitonin (detergents), we tested the subcellular protein fractionation using a digitonin-free protocol (Lodish 2000). Cells were disrupted using a 26-gauge needle and then differential centrifugation was applied as shown in Figure S2B. After analyzing the cytosolic fractions, we observed results comparable to those obtained with the digitonin-based protocol (Figure 2D and 2F). These two different protocols further strengthen the notion that ER proteins do exit the ER during stress.

Next, we sought to investigate whether the ER reflux could also apply to a large spectrum of proteins from the secretory pathway. To this end, we enriched N-glycosylated proteins from the digitonin fraction of HEK293T cells treated with Tg (Figure S2C). The purified material was subjected to mass spectrometry analysis. We focused on soluble glycoproteins that were enriched in the cytosolic fraction after Tg treatment compared to control. We identified 26 different soluble secretory N-glycoproteins present in the cytosol (Table S2). Gene Ontology-based analysis showed that most of these proteins emanated from both ER and lysosomal compartments as well as from the endomembrane system. Moreover, 23 out of these 26 proteins were part of a unique functional network (Figure 2G and Table S2), thus suggesting functional implications to this mechanism. We also performed a similar analysis on digitonin fractions from GBM tumor cells (isolated from patients) and compared them with our previous analysis of Tg treated HEK293T fractions. Interestingly, about 60% of hits were enriched in both fractionation types (Figure 2H and Table S2). This observation led us to hypothesize that under ER stress, ER protein reflux targets a broad spectrum of ER resident and secretory proteins in order to decrease the protein load within the ER.

ER protein reflux was also observed in other cancer cell lines such as GL261, U87 (GBM), and A549 (lung adenocarcinoma) (Figure 3A-H) using the two aforementioned subcellular protein fractionation protocols. Using immunofluorescence, we could colocalize the ER luminal protein PDIA1 with the cytosolic GAPDH in A549 during ER stress (Figure 4A). Moreover analysis of the immunogold electron microscopy images unveiled that there was a slight but different distribution of PDIA3 to non-ER locations in cells treated with ER stressors compared to DMSO treated cells (Figure 4B-D). Notably, in cancer cells the amount of ER proteins refluxed to the cytosol was higher than in noncancer cell lines (Figure S3A).

**Figure 3:**
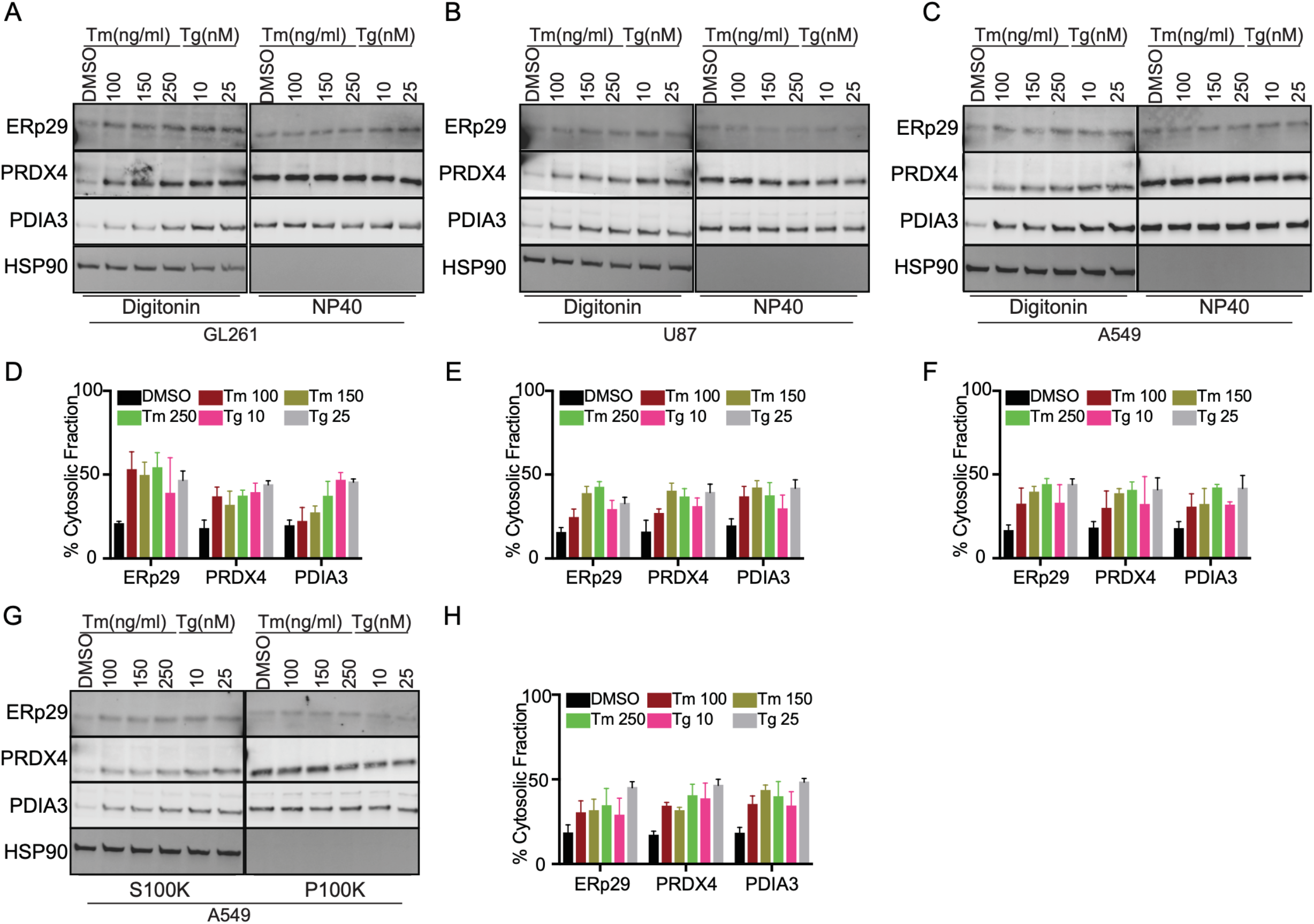
ER protein reflux is constitutive in cancer cells. (A-C) Subcellular protein fractionation of several ER resident proteins in (A) GL261, (B) U87 and (C) A549 cells treated with the indicated concentrations of Tm or Tg using Digitonin. (D-F) Quantification of the subcellular protein fractionation of several ER endogenous proteins in GL261, U87 and A549 cells as in A-C respectively. (G) Subcellular protein fractionation of several ER resident proteins in A549 cells treated with the indicated concentrations of Tm or Tg using differential centrifugation protocol. (H) Quantification of the subcellular protein fractionation of several ER endogenous proteins in A549 as in (G).

**Figure 4:**
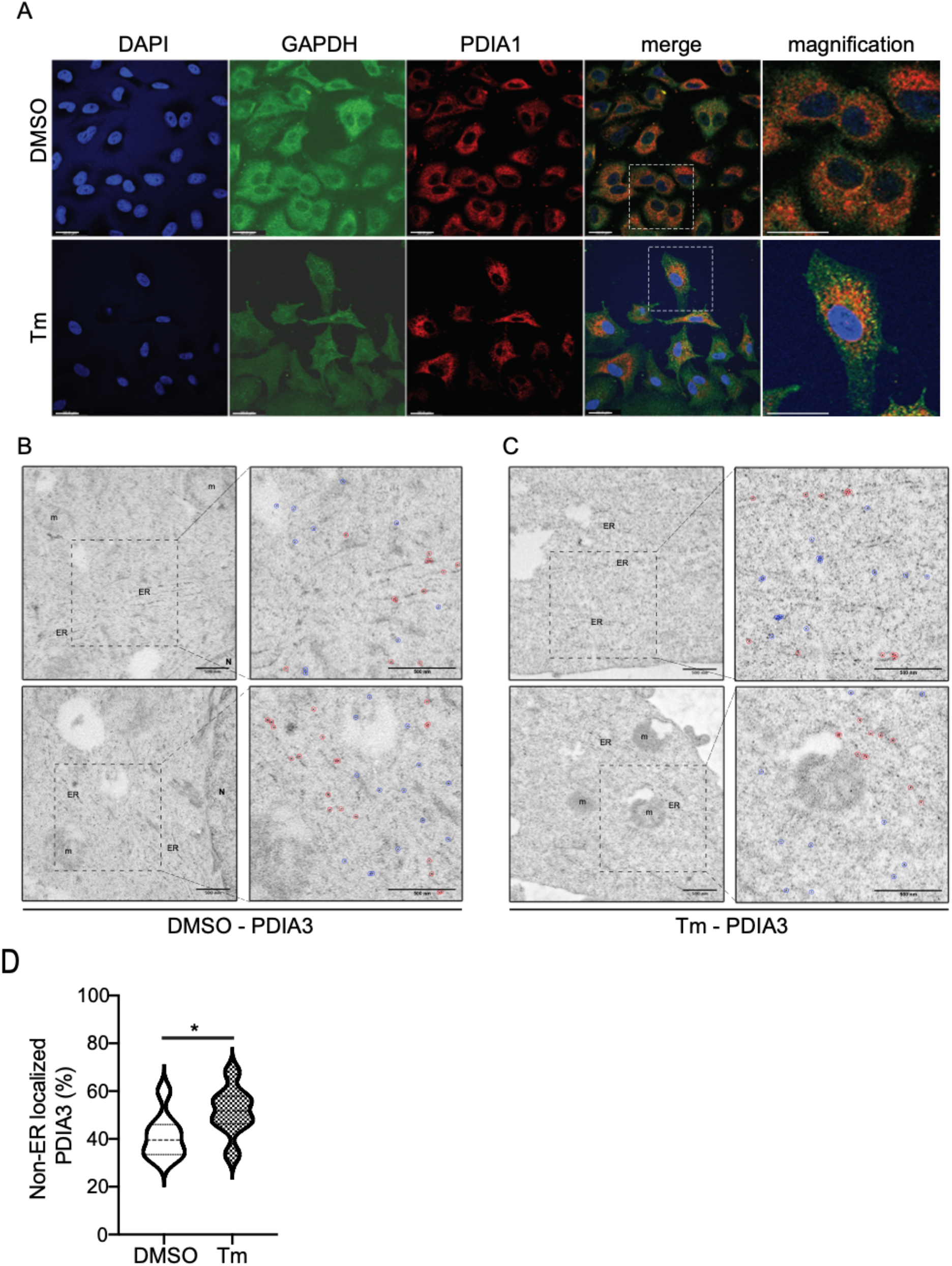
PDI proteins are redistributed to the cytosol during ER stress. (A) Immunofluorescence analysis showing colocalization between PDIA1 and GAPDH in A549 cell lines treated with DMSO or Tm for 16 hours. Cells were stained with anti-GADPH (Green) and anti-PDIA1 (Red). Nucleus were stained using Hoechst (Blue). (Scale bar 25 μm). (B-C) Representative transmission electron microscopy images of the gold particles distribution (after immunogold labelling with PDIA3 antibodies) in A549 cells treated with DMSO or Tm. In the inserts, gold particles in the ER were surrounded by red circles and those in the rest of the cytoplasm by blue circles. (Scale bar 500 nm). (D) violin plots of the gold particles distribution from the electron microscopy experiment (immunogold labelling of PDIA3) as shown in Figure 4B and 4C. *P-value=0.0486.

These data indicate that upon ER stress, ER luminal proteins (and ER targeted sfGFP/mEOS3.2) are refluxed to the cytosol. Fluorescence microscopy with ER-sfGFP and ER-mEOS3.2 in HEK293T cells in addition to the immunofluorescence experiments in A549 cells (Figure 2A-B and Figure 4) confirmed the results obtained using cell fractionation and serve as alternative detergent-free methods to monitor reflux from the ER. Moreover, the ER-mEOS3.2 experiment showed that ER protein reflux occurred for proteins that already resided in the ER rather than as a result of the pre-emptive quality control mechanism (Kang et al. 2006). ER stress mediated protein reflux may therefore act as a surveillance mechanism that is evolutionary conserved from yeast to mammals and acts as an adaptive pathway in stress conditions. This phenomenon causes ER resident proteins to reflux to the cytosol in different cell lines and was found to be constitutively active in cancer cells and in cells freshly isolated from human tumors or from murine tumor models. Moreover, the data presented herein show that this mechanism applies to a large spectrum of (glyco)proteins from the secretory pathway. Furthermore, many of the refluxed proteins identified in our experimental systems belong to a unique functional network, suggesting functional implications to this mechanism.

### ER stress mediated reflux as an ER to CYtosol Signaling (ERCYS) pathway to inhibit tumor suppressors

Thus far, we have shown that the ER protein reflux is constitutive in some cancer cells as is the activation of the UPR and we thus hypothesized that, as the UPR, it may play an adaptive and pro-oncogenic function contributing to cancer cells increased fitness. To investigate possible adaptive mechanisms of the reflux process, we evaluated the nature of refluxed proteins in A549 cells subjected to ER stress (Figure 3, Figure 5A). We focused on Anterior GRadient 2 (AGR2, PDIA17), that was highly enriched in the digitonin fractions of A549 cells during ER stress (Figure 5A). AGR2 is a PDI family member thought to catalyze protein folding through thiol-disulfide based reactions (Chevet et al. 2013). In many studies it has been shown as a proto-oncogene but how it performs this function is unclear. For instance, AGR2 was shown to inhibit the activity of the p53 tumor suppressor, but mechanism remains unknown (Pohler et al. 2004). Here, we propose a model in which ER stress in cancer cells may cause constitutive AGR2 reflux to the cytosol, where AGR2 might in turn gain new functions to interact and inhibit p53. To test this model, co-immunoprecipitation experiments showed that upon stress AGR2 was translocated to the cytosol and interacted with WT p53 in A549 cells treated with Tm, Tg and BFA (Figure 5B). We documented this mechanism by measuring wild-type (wt) p53 transcriptional and p21 protein expression levels (a downstream target of p53 signaling). Tm, Tg and BFA treatment reduced p21 protein levels, as well as wt-p53 phosphorylation and transcriptional activity as shown in cells transfected with a luciferase reporter under the p53-DNA binding site (Figure 5C). Moreover, AGR2-silenced A549 cells showed increased p53 phosphorylation and p21 protein levels under ER stress conditions compared to control cells (Figure 5D). This confirms that AGR2 is involved in the inhibition of wt-p53 activity under ER stress. To further document that the observed inhibition of wt-p53 is linked to the presence of refluxed AGR2, we engineered two nanobodies to specifically target AGR2 (either in the ER or in the cytosol) (Figure 5E). Notably, both AGR2 nanobodies showed minimal decrease in p21 protein levels and p53 phosphorylation compared to cells transfected with control nanobodies (Figure 5E). These data show that cytosolically localized AGR2 gains new functions through interacting and inhibiting wt-p53 activity. As such, targeting cytosolic AGR2 in cancers could be used to restore p53 pro-apoptotic transcriptional activity and sensitize them to existing anti-cancer therapies. Alternatively, in pre-neoplastic stages such as Barrett’s oesophagus, in which AGR2 was first described to inhibit p53 (Pohler et al. 2004), the targeting of cytosolic AGR2 might prevent the inhibition of p53 tumor suppressor activity, thereby lowering their transformation potential.

**Figure 5:**
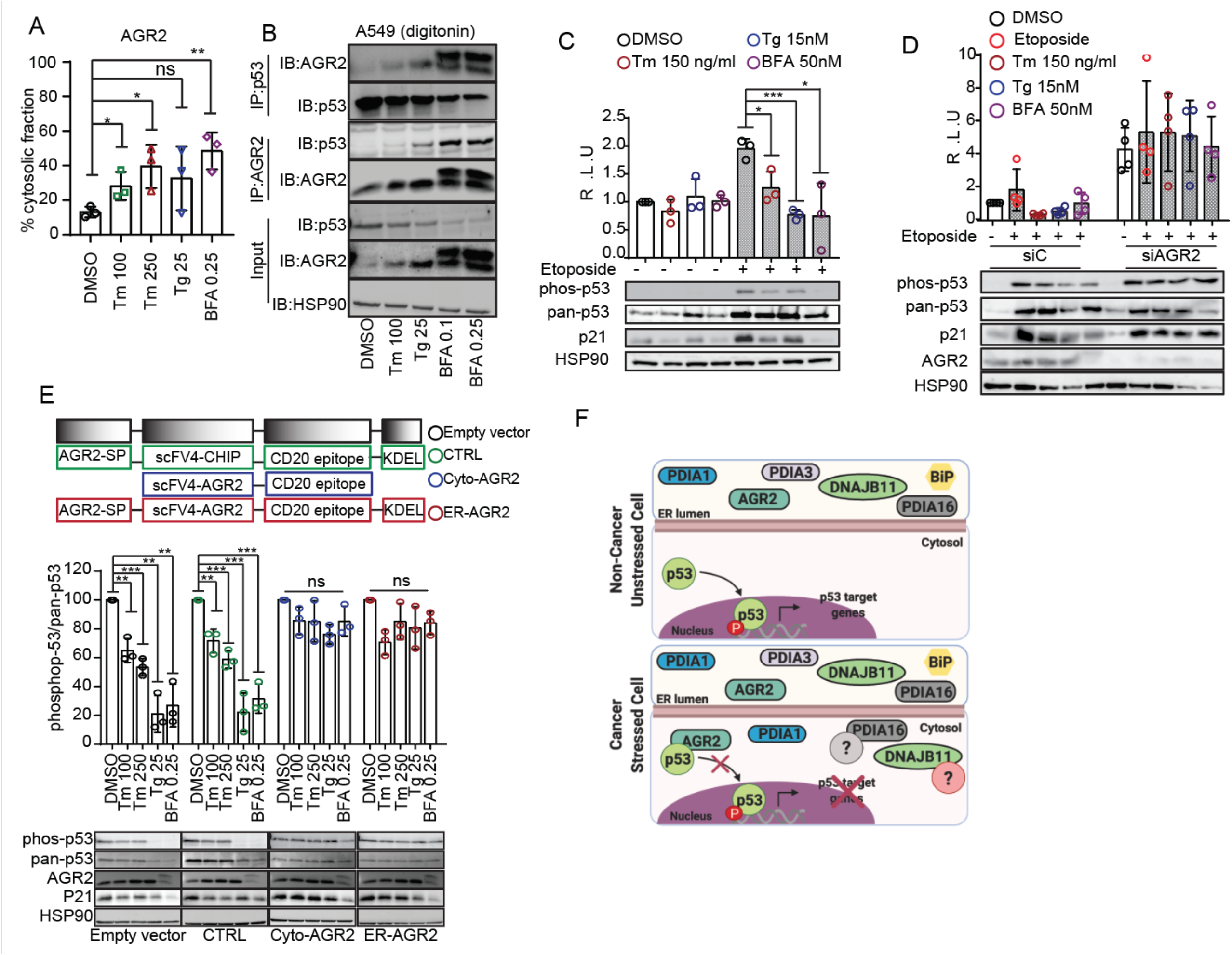
AGR2 reflux from the ER to the cytosol in A549 cells results in non-genetic inactivation of p53. (A) Quantification of the subcellular protein fractionation of AGR2 in A549 cells treated with Tm, Tg and BFA for 16 hrs. (B) Immunoprecipitation of p53 and AGR2 in the digitonin fraction of A549 cells. (C) A549 were treated with Tm, Tg and BFA at the indicated concentrations for 16 hours. Luciferase experiments were performed after 24 hours of transfection. Graph shows the fold induction of p53 luciferase construct. Western blot experiments for phospho-p53, pan-p53, p21 and HSP90 were performed as control. (D) A549 were transfected with control SìRNA (siC) and AGR2-tergetd siRNA (siAGR2). After 24 hours, cells were transfected with p53-luciferase construct. Cells were then treated as in C and luciferase experiments were performed. (E) A549 were transfected with the indicated constructs of differently targeted nanobodies, cells were then treated with Tm, Tg and BFA for 16 hours at the indicated concentration. Western blot experiments were performed and HSP90 were used as loading control. (F) Cartoon showing our working model, ER stress induction promotes reflux of some PDI and PDI-L proteins including AGR2 to the cytosol. In the cytosol, AGR2 is able to bind and inhibit p53 function.

## DISCUSSION

The phenomenon described in this manuscript, named ER-to-CYtosol-Signaling (ERCYS), may also play an important physiological role. To date several mechanisms were reported to decrease ER protein load either through signaling mechanisms including regulated IRE1-dependent decay (RIDD) of RNA, PERK-mediated global protein translation attenuation or through clearance mechanisms including pre-emptive quality control and ERAD processes. However, here, we demonstrate that ERCYS can reduce secretory protein load in the ER lumen by refluxing folded and mature proteins to the cytosol during stress. Moreover, ERCYS-mediated protein reflux into the cytosol is associated with a selective gain-of-function that is pivotal for cancer development. Indeed, this mechanism as illustrated by the cytosolic gain-of-function acquired by AGR2, could represent a unique non-genetic process of tumor suppression through an interaction with p53. As such this could be one of many other mechanisms by which the refluxed ER proteins in tumor cells could influence existing proapoptotic signaling pathways in the cytosol (Figure 5F). Further studies on the role(s) of the cytosolically localized AGR2 and other PDI-like proteins as well as on the precise nature of the refluxed ER proteins will certainly impact on the understanding ER stress mediated diseases including cancer and will open new areas for biological exploration and therapeutic strategies.

## MATERIALS AND METHODS

### Cell culture and transfection

Human HEK293T, A549, MCF7, U87 and mouse GL261 cell lines cells were cultured in Dulbecco’s modified Eagle’s medium (DMEM) supplemented with 10% FBS and 1% antibiotics at 37°C in a 5% CO2 incubator. PG13-luc (WT p53 binding sites) was a gift from Bert Vogelstein (Addgene plasmid # 16442; http://n2t.net/addgene:16442; RRID: Addgene_16442). Cells were transfected with Lipofectamine LTX and plus reagent or Lipofectamine 2000 (Thermo-Fisher Scientific) according to the manufacturer’s protocols. Small-interfering RNAs (siRNA) were obtained from Ambion Each siRNA (25 nM) was transfected by reverse transfection using Lipofectamine RNAiMAX (Invitrogen). Thapsigargin (Tg), Etoposide (Eto), Brefeldin A (BFA) were obtained from Sigma-Aldrich (St. Louis, MO, USA). Tunicamycin (Tm) was purchased from Calbiochem.

### Immunoblot and immunoprecipitation

Whole cell extracts were prepared using RIPA buffer (25 mM Tris/HCl pH 7.5, 150 mM NaCl, 1% NP-40, 1% sodium deoxycholate, and 0.1% SDS). Primary antibodies were incubated overnight at 4°C. Secondary antibodies were incubated 1h at RT (1:7000), the antibodies used in this work are listed in Table S3. For the immunoprecipitation analysis, cells were lysed in Co-IP buffer (50 mM Tris/Hcl pH 8, 150 mM NaCl, 0.5% TritonX100 and 1 mM EDTA), incubated 30’ on ice and then incubated 16 h at 4°C with the anti-AGR2 or anti-p53 antibodies (1 μg Ab/1000 μg protein). After this, Dynabeads protein G/A (Life Technologies) were first washed with CoIP lysis buffer, then mixed with the protein/Ab mixture, incubated at 4°C for 3 hours with gentle rotation and washed with Co-IP buffer. Finally, the beads were eluted with 50 μl of Laemmli sample buffer, heated at 100°C for 5 min and loaded to SDS/PAGE. For the immunoblotting, anti-AGR2 or anti-p53 antibodies were used.

### Tumor isolation

Mouse and human brain samples were collected and mechanically dissociated using gentleMACS dissociation following the manufacturer’s instructions (Miltenyi Biotec, Paris, France). Freshly isolated tumor tissues were suspended in 5ml DMEM immediately after being surgically removed and placed in a petri dish. Tumor and non-tumor tissues were cut to small pieces of 1-2 mm^3^ using a sterile scalpel and then transferred to C-tube and tightly closed. Then we used the mechanical dissociation program A.01 for C-tubes (the most gentle program to homogenize tissues). The resulting homogenate was then directly decanted into 40 μm cell strainer into 50ml tube. Cells were then pelleted at 100 RCF for 5 minutes washed and re-pelleted for another 5 minutes before being subjected to 25 μg/mL digitonin as shown below in the subcellular protein fractionation protocol.

### Differential centrifugation

Cells were washed with PBS and trypsinized for 5minutes, pellets were collected at 300 x g and washed again with ice cold PBS. Cells were suspended in homogenizing buffer (20 mM HEPES pH7.4, 10 mM KCl, 2 mM MgCl2, 1 mM EDTA, 1 mM EGTA, 1 mM DTT and protease inhibitor cocktail) and then passed 15 times through a 26-gauge needle (1 mL syringe). After incubation for 20 minutes on ice, the lysates were centrifuged 300 RCF for 5 minutes at 4°C. The pellet (P300) contained cell debris and nuclei while the supernatant (S300) contained cytoplasm, membranes and mitochondria. S300 then was further centrifuged for 5 minutes at 13,000 RCF, the pellet (P13,000) contained the mitochondria and the supernatant (S13,000) contained the cytoplasm and the membrane fraction. S13,000 then was centrifuged for another 1 hour at 100,000 RCF in an ultracentrifuge. After recovering the supernatant (S100K-Cytoplasmic Fraction), the pellet (P100K-membrane fraction) was resuspended in 400 μL of Laemmli sample buffer.

### Digitonin permeabilization

To obtain cytosolic and membrane fractions, cultured cells and dissociated cells from mice and human brains were subjected to subcellular fraction as shown in (Holden and Horton 2009). In brief, cells were washed with cold PBS and trypsinized for 5’, collected and pellets were obtained after centrifugation at 100 RCF for 5’. Pellets were washed twice with PBS and then resuspended in Buffer 1 (see below). After 10’ in gently rotation at 4°C, tubes were subject to centrifugation at 2000 RCF at 4°C for 5’ and supernatants were harvested (Fraction 1-cytosol). The obtained pellets were then resuspended in Buffer 2 and incubated 30’ on ice. After 10’ centrifugation at 7500 RCF at 4°C, supernatants were harvested a (Fraction 2-membranes). Buffer 1 (cytosol) – 50 mM HEPES pH 7.4, 150 mM NaCl, 10 μg/ml digitonin (add fresh). Buffer 2 (membranes) – 50 mM HEPES pH 7.4, 150 mM NaCl, 1% NP40.

### Mass spectrometry analysis

*Protein digestion: After protein precipitation with acetone*, pellet was denatured with 8 M urea in 0.1 M Tris-HCl pH 8.5, reduced at 50°C with 5 mM TCEP for 30 min and alkylated with 10 mM iodoacetamide for 30 minutes in the dark. First digestion was performed with 5 μg endoproteinase Lys-C (Wako) urea for 6h, followed by a 4 times dilution in Tris buffer and an overnight trypsin digestion (Promega) at a ratio 1/100. Digestion was stopped with formic acid (1% final) and peptides were desalted on Sep Pak C18 cartridge (Waters Corporation). Peptides dissolved in 0.1% TFA were quantified using colorimetric assay (Pierce - ThermoFisher Scientific) and adjusted at 5 mg/ml. *N-glycopeptide enrichment:* N-glycopeptide enrichment is based on a protocol previously described (Yakkioui Y et al, 2017). Peptides (500 μg in 0.1 M sodium acetate pH 5.5, 150 mM NaCl) were incubated with NaIO4 for 1 h then 15 min with NaS2O3 and mixed with 100 μl AffiGel Hz Hydrazide Gel (Biorad) overnight. The unbound material was washed with tris 0.1M pH 8.5, glycine 0.1M, isopropanol 10% then 3 times with PBS. Lastly, PNGase F was added for 6h at 37°C and the deglycosylated peptides were extracted with 0.5% TFA. *LC-MS analysis:* Total-fraction (i.e. not Nglyco enriched) and glyco-fraction were analyzed with an Orbitrap ELITE coupled with a nanoLC chromatographic system (ThermoFisher Scientific). Briefly, peptides were separated on a C18 nano-column with a linear gradient of acetonitrile and analyzed with a Top 20 CID method. Each sample was analyzed in triplicate. Data were processed by database searching against Human Uniprot Proteome database using Proteome Discoverer 2.3 software (Thermo Fisher Scientific). Precursor and fragment mass tolerance were set at 10 ppm and 0.6 Da respectively. Trypsin with up to 2 missed cleavages was set as enzyme. Oxidation (M, +15.995 Da), Deamidation (N, +0.984) were set as variable modification and Carbamidomethylation (C, + 57.021) as fixed modification. Peptides and proteins were filtered with False Discovery Rate <1%. N-glycopeptides were filtered based on the detection of deamidation and the presence of the consensus motif NxS/T. Lastly quantitative values obtained from Extracted Ion Chromatogram (XIC) were exported in Perseus for statistical analysis.

### Immunoelectron microscopy

Cell cultures were fixed in a 2.5% PFA, 0.05% glutaraldehyde solution in 80 mM Sorensen phosphate buffer pH 7.4 for 1 h at 4°C and were scraped and washed 15 min with 0.1 M phosphate buffer. To facilitate handling of the cells, they were coated with 1.5% agarose and then cut into 1 mm3 pieces. Cells were dehydrated in ethanol (30–100%) on ice and gradually infiltrated with ethanol/LR White resin (Delta microscopies) (2/1; 1/1; 1/2; successively) and finally infiltrated with pure resin overnight at 0°C. Polymerization was carried out at 60°C for 24h. Thin sections (80 nm) were collected onto 300 mesh nickel grids, and processed for immunochemistry. Sections were blocked in 20mM Tris–HCl pH 7.6, 150mM NaCl (TBS buffer), containing 1% BSA, 0.1% BSA-c™ (Aurion), 10% goat serum (Aurion), 0.2% Tween 20, twice 20 min. Grids were then incubated for 2h at room temperature with anti-PDIA3 rabbit antibody (dilution 1:25) in Ab-buffer (TBS pH 7.6, 1% BSA, 0.1% BSA-c™, 1% goat serum, 0.2% Tween 20). Following four washes with the Ab-buffer, grids were incubated for 1 h with goat antirabbit Ig conjugated to 10 nm colloidal gold (1:40 dilution) in Ab-buffer. Sections were washed in Ab-buffer, fixed in 2.5% glutaraldehyde, and finally stained with 5% uranyl acetate and lead citrate. Negative controls were carried out, omitting primary antibodies. Sections were examined on a Tecnai Sphera operating at 200 kV (FEI, Eindhoven, Netherlands), and images were recorded with a 4×4 k CCD Ultrascan camera (Gatan, Pleasanton, USA). We used at least 8 images (x 14,500) displaying both ER and cytoplasm for each condition. The distribution of PDIA3 was estimated by calculating the ratio between the particle in the ER and the rest of the cytoplasm.

### Mouse work

*Tumor cell orthotopic Implantation* – Tumor cells (GL261) were implanted in the brain of immunocompetent C57BL/6rJ, 8 weeks old male mice *(Janvier, Laval, France) and tumor cells (U87)* were implanted in the brain of immunodeficient mice NSG *(NOD.Cg-Prkdcscid Il2rgtm1Wjl/SzJ)* mice were purchased from Charles River Laboratories (Wilmington, MA), 8 weeks old male mice *(Janvier, Laval, France)*. All animal procedures met the European Community Directive guidelines (Agreement B35-238-40 Biosit Rennes, France/ No DIR 13480) and were approved by the local ethics committee and ensuring the breeding and the daily monitoring of the animals in the best conditions of well-being according to the law and the rule of 3R (Reduce-Refine-Replace). GL261-Luc cells were implanted in the mouse brain by intracerebral injection followed by tumor growth analysis using bioluminescence. The mice were anesthetized intraperitoneally and then fixed on a stereotactic frame. After incising the scalp, the stereotaxic coordinates were calculated for injection of tumor cells into a specific point of the brain, and reproducible for all the mice used. In the study, the tumor cells, 2,5.10^4^ cells per mice in 1 μL for GL261-luc and 5×10^4^ cells per mice in 1 μL for U87, are injected at 2.2 mm to the left of the Bregma and 3.2 mm deep to perform the implantation at the level of the striatum.

### Immunofluorescence

Cells were fixed with paraformaldehyde (4%) at room temperature for 15 minutes, permeabilized with PBS containing 0.1% TritonX100 for 10 minutes and blocked with PBS containing 5% BSA for 30 minutes. Fixed and permeabilized cells were then incubated for 1hour with primary antibodies (1:100 dilution) in PBS + 0.1% Saponin. Slides were washed three times with PBS and incubated with goat anti-mouse IgG (H+L) Alexa Fluor^®^ 647 (ThermoFisher #A21235) and goat antimouse IgG (H+L) Alexa Fluor^®^ 488 (ThermoFisher #A11001) conjugated secondary antibodies for 30 minutes at 37°C. Images were captured using a Leica SP8 confocal microscope.

## ACKNOWLEDGEMENTS

We thank Giannino Del Sal (LNCIB, Trieste, Italy) for anti-p53 (Valentino) antibodiy. We also thank David Y Thomas (McGill University, Canada) and Gisou van der Goot (EPFL, Switzerland) for critically reading the manuscript. This work was funded by grants from Institut National de la Santé et de la Recherche Médicale (INSERM), Institut National du Cancer (INCa, PLBIO), Fondation pour la Recherche Médicale (FRM, équipe labellisée 2018), Agence National de la Recherche (ANR, ERANET ERAAT), MSCA RISE-734749 (INSPIRED) grants to EC; the BBSRC UK (EM; BB/C511599/1; and BB/J00751X/1) and UKM Fellowship and Ministry of Higher Education (MOHE) of Malaysia KPT(BS)870809015063 (MAM). DS was supported by an AIRC fellowship for Abroad. This work has benefited from the Microscopy Rennes Imaging Center (Mric) facility for electron microscopy.

