## Supplementary Figures and tables for "Reflux of Endoplasmic Reticulum proteins to the cytosol yields inactivation of tumor suppressors"

**Supplementary tables** Table S1 – S3

**Supplementary figure legends** Figure S1 – S3

**Supplementary figures** Figure S1 – S3

### SUPPLEMENTARY TABLES

#### **Table S1**: Characteristic of human patients involved in this study

| Patient | Age at diagnosis | Gender | Symptoms | Karnofsky index | Location | IDH status | ATRX expression |
|---|---|---|---|---|---|---|---|
| 1 | 72 | Male | Aphasia | 80 | Left frontal lobe | WT | conserved |
| 2 | 72 | Male | Memory disturbance | 70 | Right temporal lobe | Mutated | conserved |
| 3 | 65 | Female | Cognitive disturbance | 70 | Corpus callosum | WT | conserved |
| 4 | 70 | Female | Seizures | 100 | Right frontal lobe | Mutated | conserved |
| 5 | 70 | Male | Cognitive disturbance | 80 | Left temporal lobe | WT | conserved |
| 6 | 53 | Male | Headache | 90 | Right temporal lobe | WT | conserved |
| 7 | 73 | Male | Cognitive disturbance | 70 | Right temporal lobe | WT | conserved |
| 8 | 50 | Male | Seizures | 90 | Right frontal lobe | WT | conserved |
| 9 | 67 | Female | Seizures, motor deficit | 70 | Left frontal lobe | WT | conserved |

#### **Table S2**: List of the identified N-glycopeptides from the HEK293T cells treated with Tg and from the cytosolic fraction of isolated human GBM tumors.

| Accession | Description | split after | baseGlyco | target | Coverage [%] | # Peptides | # PSMs | Jnique Peptic | # AAs | MW [kDa] |
|---|---|---|---|---|---|---|---|---|---|---|
| P14625 | Endoplasmin OS=Homo sapiens OX=9606 | HSP90B1 | + | + | 54 | 44 | 1063 | 43 | 803 | 92,4 |
| Q9NYU2 | UDP-glucose:glycoprotein glucosyltransferase 1 OS=Homo sapiens OX=9606 | UGGT1 | + | + | 22 | 25 | 173 | 25 | 1555 | 177,1 |
| Q96HE7 | ERO1-like protein alpha OS=Homo sapiens OX=9606 | ERO1A | + | + | 41 | 15 | 104 | 14 | 468 | 54,4 |
| Q08380 | Galectin-3-binding protein OS=Homo sapiens OX=9606 | LGALS3BP | + | + | 25 | 9 | 78 | 9 | 585 | 65,3 |
| P11047 | Laminin subunit gamma-1 OS=Homo sapiens OX=9606 | LAMC1 | + | + | 19 | 20 | 66 | 20 | 1609 | 177,5 |
| Q9H3G5 | Probable serine carboxypeptidase CPVL OS=Homo sapiens OX=9606 | CPVL | | + | 27 | 10 | 60 | 10 | 476 | 54,1 |
| P53634 | Dipeptidyl peptidase 1 OS=Homo sapiens OX=9606 | CTSC | + | + | 21 | 6 | 59 | 6 | 463 | 51,8 |
| P13674 | Prolyl 4-hydroxylase subunit alpha-1 OS=Homo sapiens OX=9606 | P4HA1 | + | + | 25 | 10 | 58 | 10 | 534 | 61 |
| Q96AY3 | Peptidyl-prolyl cis-trans isomerase FKBP10 OS=Homo sapiens OX=9606 | FKBP10 | + | + | 24 | 9 | 47 | 9 | 582 | 64,2 |
| Q8NBJ5 | Procollagen galactosyltransferase 1 OS=Homo sapiens OX=9606 | COLGALT1 | | + | 15 | 9 | 43 | 9 | 622 | 71,6 |
| Q9UHG3 | Prenylcysteine oxidase 1 OS=Homo sapiens OX=9606 | PCYOX1 | + | + | 17 | 8 | 41 | 8 | 505 | 56,6 |
| P10253 | Lysosomal alpha-glucosidase OS=Homo sapiens OX=9606 | GAA | | + | 17 | 9 | 31 | 9 | 952 | 105,3 |
| Q99538 | Legumain OS=Homo sapiens OX=9606 | LGMN | + | + | 17 | 4 | 28 | 4 | 433 | 49,4 |
| Q8NHP8 | Putative phospholipase B-like 2 OS=Homo sapiens OX=9606 | PLBD2 | + | + | 12 | 6 | 18 | 6 | 589 | 65,4 |
| P08236 | Beta-glucuronidase OS=Homo sapiens OX=9606 | GUSB | | + | 8 | 4 | 12 | 4 | 651 | 74,7 |
| Q12841 | Follistatin-related protein 1 OS=Homo sapiens OX=9606 | FSTL1 | | + | 13 | 4 | 10 | 4 | 308 | 35 |
| P07711 | Cathepsin L1 OS=Homo sapiens OX=9606 | CTSL | | + | 8 | 2 | 7 | 2 | 333 | 37,5 |
| Q9H497 | Torsin-3A OS=Homo sapiens OX=9606 | TOR3A | + | + | 7 | 2 | 5 | 2 | 397 | 46,2 |

**Table S3**: List of the antibodies used in this study

| Antibody | Company |
|---|---|
| IRE1$\alpha$ | CST/3294S |
| PERK (C33E10) | CST/3192S |
| ATF6 (1-7) | Abcam/ab122897 |
| PDIA3 (Mouse) | Ptg/66423-1-Ig |
| PDIA3 (Rabbit) | Ptg/15967-1-AP |
| PDIA16 | Abcam/ab134938 |
| PDIA9 | Ptg/24344-1-AP |
| PDIA1 (Mouse) | Ptg/66422-1-Ig |
| PDIA1 (Rabbit) | Ptg/11245-1-AP |
| AGR2 (Rabbit) | Ptg/12275-1-AP |
| AGR2 (Mouse) | SantaCruz/sc-101211 |
| DNAJB11 | Ptg/15484-1-AP |
| PRDX4 | Ptg/10703-1-AP |
| pan-p53 (DO-1) (Mouse) | SantaCruz/sc-126 |
| pan-p53 (Valentino) (Rabbit) | Gift from LNCIB(Girardini et al. 2011) |
| phospho-p53 (Ser15) | Cell signaling #9284 |
| p21 Waf1/Cip1 (12D1) | Cell signaling #2947 |
| Calnexin | Gift from JJM Bergeron (McGill, Canada) (Ou et al. 1993) |
| GADPH (G-9) | SantaCruz/sc-365062 |
| HSP90 (4F10) | SantaCruz/sc- 69703 |

### SUPPLEMENTARY FIGURES LEGENDS

**Figure S1. ER protein reflux in human and mouse derived GBM tumors.** (A) schematic of the subcellular protein fractionation experiment flow. (B) Western blot for the UPR markers in total cell lysates from mice-derived tumor and non-tumor tissues. (C) Western blot experiments of ER-lumenal proteins of U87-grafted-mice-derived tumor masses. Lower panel show percentage of proteins detected in the cytosolic fraction. (D) Schematic of our workflow for GBM Human-patient derived tumors.

**Figure S2: Schematics for the different protocols used in this study.** (A) Schematic representation of the ER-targeted yemEos3.2 construct and principles of function. (B) Flowchart representation of the differential centrifugation protocol used in this study to obtain the cytosolic fraction (C) Proteomics experiments workflow. Right panel show Volcano plot before and after N-Glyco protein enrichment.

**Figure S3: ER protein reflux is constitutive in cancer cells.** (A) Graph show the percentage of ER-lumenal proteins in the cytosol of non -cancer (MRC5, MEF and HEK293T) and cancer cells (A549, MCF7 and GL-261) in non-stressed conditions.

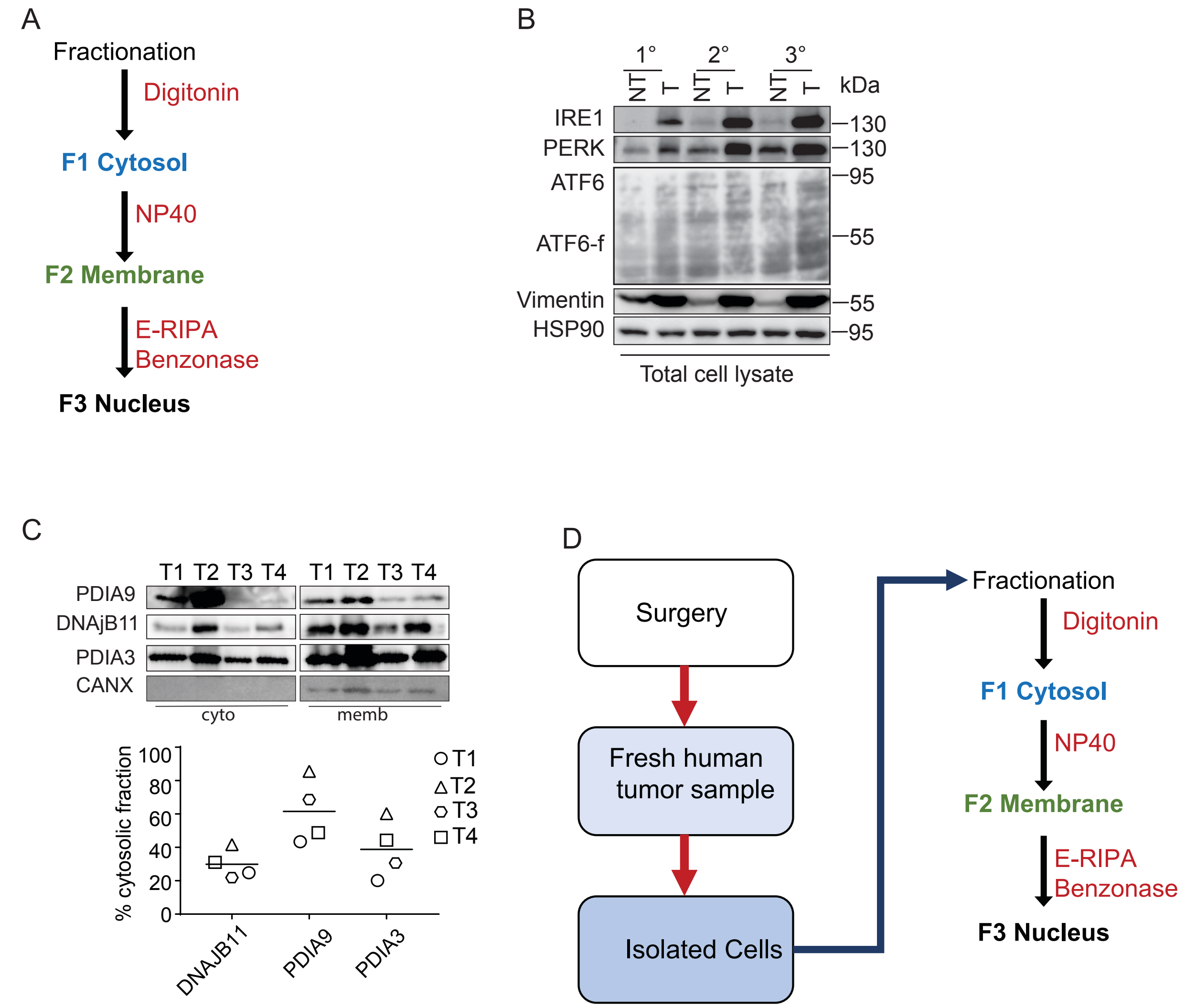


Sicari et al, Figure S1

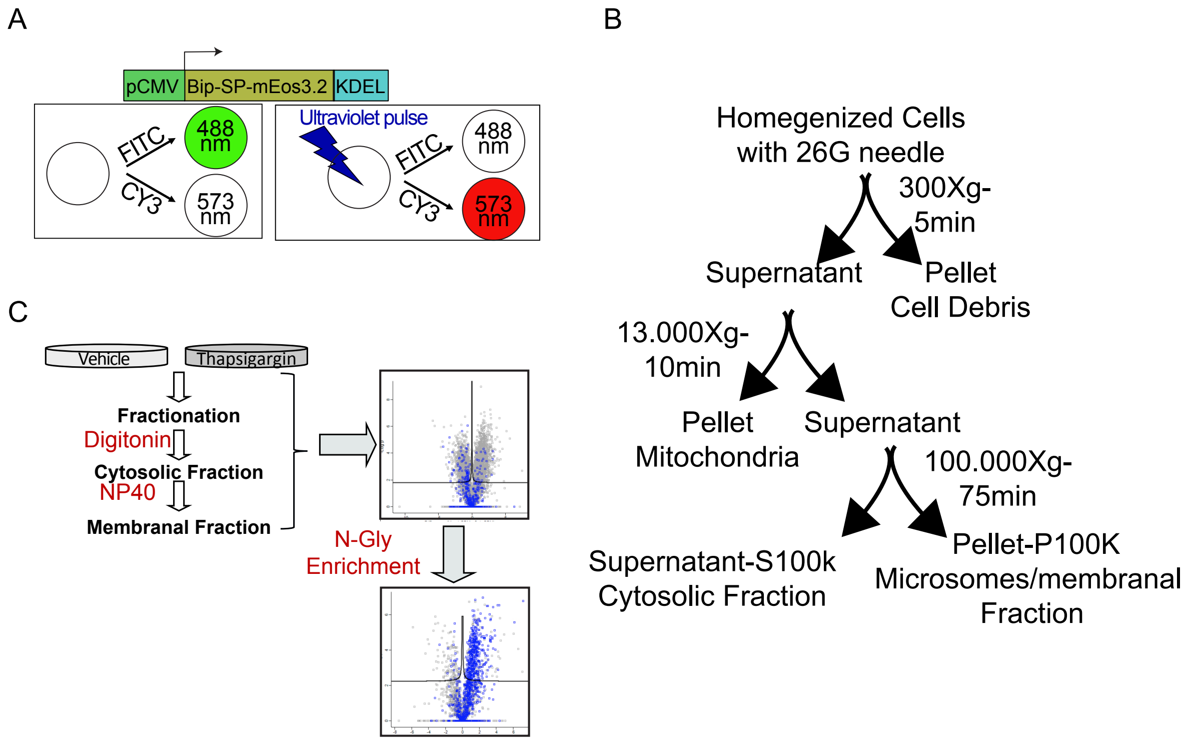


Sicari et al, Figure S2

A

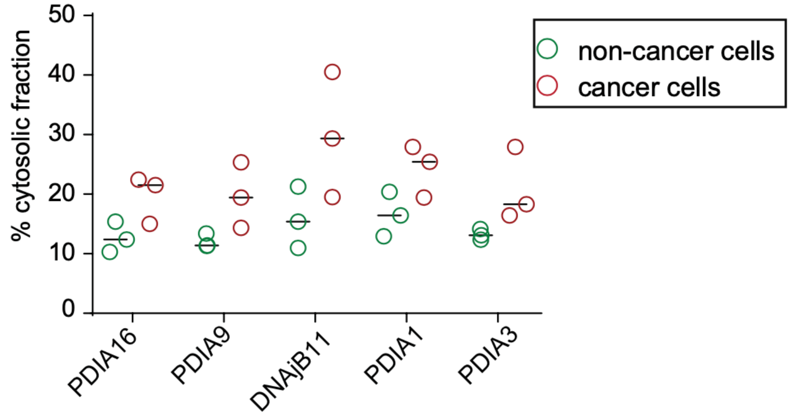


Sicari et al, Figure S3
